# Fine-Tuning Transformers For Genomic Tasks

**DOI:** 10.1101/2022.02.07.479412

**Authors:** Vlastimil Martinek, David Cechak, Katarina Gresova, Panagiotis Alexiou, Petr Simecek

**Author notes:** Correspondence to: Petr Simecek < >. Equal contribution (Vlastimil Martinek, David Cechak).

## Abstract

Transformers are a type of neural network architecture that has been successfully used to achieve state-of-the-art performance in numerous natural language processing tasks. However, what about DNA, the language life written in the four-letter alphabet? In this paper, we review the current state of Transformers usage in genomics and molecular biology in general, introduce a collection of benchmark datasets for the classification of genomic sequences, and compare the performance of several model architectures on those benchmarks, including a BERT-like model for DNA sequences DNABERT as implemented in HuggingFace (armheb/DNA_bert_6 model). In particular, we explore the effect of pre-training on a large DNA corpus vs training from scratch (with randomized weights). The results presented here can be used for identification of functional elements in human and other genomes.

## 1. Introduction

In the past five years, Deep Learning methods for Natural Language Processing (NLP) came through a revolution that has been possible thanks to two key novel innovations: language models and transfer learning. With this approach, the model is first trained in an unsupervised fashion with unlabelled data and then fine-tuned to a specific downstream task with labelled data. (Howard & Ruder, 2018) trained the ULMFit model to predict the following word in English Wikipedia corpus and then fine-tuned it to six text classification tasks (outperforming the state-of-the-art methods at a time). While ULMFit architecture was still based on Long Short Term Memory networks (LSTMs), the novel model architecture based on Encoder / Decoder structure and self-attention was introduced at around the same time – Transformers (Vaswani et al., 2017) – and have dominated the NLP field since then. It was shown that some neurons and attention heads have a direct connection to text features like sentiment (Radford et al., 2017) or direct objects of verbs (Clark et al., 2019). While the original transformer models like BERT (Devlin et al., 2018) have just lower hundred millions of parameter, much larger language models have been recently introduced like GPT-3 with 175B parameters (Brown et al., 2020), Gopher with 280B parameters (Rae et al., 2021) and GLaM with more than 1.2T parameters (Du et al., 2021). Other recent changes include the unification of different tasks (Raffel et al., 2019) expansion of transformer architecture beyond traditional sequential models, e.g. Vision transformers (Dosovitskiy et al., 2020), (Dai et al., 2021) and/or 3D Point Cloud transformers (Zhao et al., 2020).

But what about DNA, the language life written in the four-letter alphabet? For the simplicity reasons, we restrict ourselves to the human genome in this paper. It consists of more than 3 billion base pairs organized into 22 paired chromosomes (autosomes) and the 23rd pair of sex chromosomes (XX for females, XY for males). The known successful deep learning applications for convolutional neural networks (CNNs) and recurrent LSTMs include identification/classification of genes from their sequence (Georgakilas et al., 2019) and identification of functional elements regulating gene expression, namely gene promoters (Umarov & Solovyev, 2017), enhancers (Liu et al., 2016), enhancer-promoter interactions (Zeng et al., 2018) and transcription factor binding sites (Shen et al., 2018).

Unfortunately, unlike in NLP, there are no widely recognized DNA benchmarks. To overcome this problem, we have started to work on a collection of genomic datasets and propose the first five of them in the Method section. The second issue is more serious, DNA is written rather in several languages than one original language. The ∼ 20, 000 protein coding gene sequences represent ∼ 1% of the human genome. Approximately 50% of the human genome is made up of repetitive sequences, mostly transposons, but also microsatellites and minisatellites and even duplications of large segments (Haubold & Wiehe, 2006).

There are also not so many language models trained for DNA. (Hoarfrost et al., 2020) trained ULMFit-like model LookingGlass on microbial genomes. Karl Heyer published his experiments as GitHub repo GenomicULMFit (https://github.com/kheyer/Genomic-ULMFiT). Regarding transformer architecture, to the best of our knowledge, the only known language model is DNABert (Ji et al., 2020) trained on the human genome that we will utilise for our purposes. While it is not explicitly mentioned inDNABert paper, the model can be found in HuggingFace model repository (armheb/DNA_bert_6_model).

## 2. Methods

### 2.1. Datasets

Due to the lack of established genomic benchmarks, we have started to put together our own. The collection is based on a combination of existing datasets obtained from published papers and novel datasets constructed from public databases. The data are distributed as a Python package available at https://github.com/ML-Bioinfo-CEITEC/genomic_benchmarks, the minimalist version (compressed list of genomic coordinates) is stored on GitHub itself and full datasets (full DNA sequences) are cached on Google Drive.

For this paper, we have used the five datasets that have already been curated and will be part of the benchmark in the future. For testing we use ∼ 30% of data points. All five datasets contains exactly two classes and are either balanced or (in case of human promoters) close to it. The summary table with number of sequences and their lengths are in Table 1.

**Table 1.**
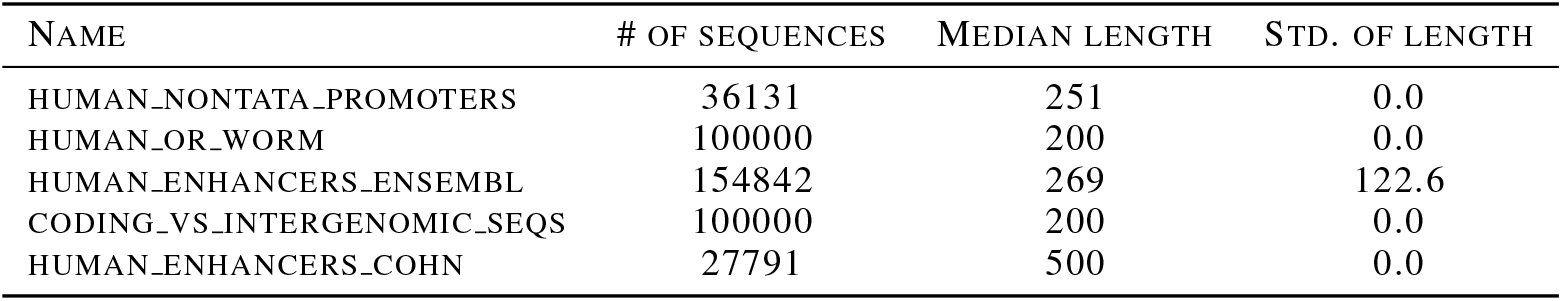
Number of sequences and sequence length per dataset.

| NAME | # OF SEQUENCES | MEDIAN LENGTH | STD. OF LENGTH |
| --- | --- | --- | --- |
| HUMAN_NONTATA_PROMOTERS | 36131 | 251 | 0.0 |
| HUMAN_OR_WORM | 100000 | 200 | 0.0 |
| HUMAN_ENHANCERS_ENSEMBL | 154842 | 269 | 122.6 |
| CODING_VS_INTERGENOMIC_SEQS | 100000 | 200 | 0.0 |
| HUMAN_ENHANCERS_COHN | 27791 | 500 | 0.0 |

#### 2.1.1. Human Non-TATA Promoters

A promoter is a sequence of DNA that binds a protein initiating the gene transcription. Effectively, it turns gene expression on and off. It is usually located close (from -200 to 50bp) to the transcription splice site (TSS). This dataset has been adapted from the paper (Umarov & Solovyev, 2017).

#### 2.1.2. Human Enhancers Cohn

An enhancer is a sequence of DNA that can bound specific proteins and therefore increase a change of transcription of a particular gene. Unlike promoters, enhancers do not need to be in a close proximity to TSS (might be several Mb away). This dataset has been adapted from (Cohn et al., 2018) paper.

#### 2.1.3. Human Enhancers Ensembl

For this dataset of human enhancers, we have queried Ensembl database (Howe et al., 2021), release 100. The data are originally coming from VISTA Enhancer Browser project, (Visel et al., 2007). The Unlike the other datasets, this one has variable length of the sequences.

#### 2.1.4. Coding Vs Intergenomic Regions

This dataset has been originally used for teaching purposes at ECCB2020 workshop. It consists of randomly generated 50,000 sequences (200bp long) from intergenomic regions and randomly generated 50,000 sequences from human transcripts.

#### 2.1.5. Human Or Worm?

Randomly chosen DNA sequences (200bp long) either from the human genome or from the genome of C. elegans (worm).

### 2.2. Models & Training

We have trained and evaluated three models for each dataset: First, we fine-tuned DNABert model pre-trained on human DNA (Ji et al., 2020). Second, to assess the effect of pre-training, we trained the model initialized with random weights (no pre-training). Lastly, as a baseline we have then used CNN architecture previously successfully used to similar problems (Klimentova et al., 2020).

We have repeated each training five time to evaluate the variability of the results. As a loss function, we have used binary cross entropy. We have used early stopping and fallback to the model that achieved the lowest loss on the validation set.

The BERT models were trained with batch size of 48 and weight decay of 0.1. The learning rate was linearly increased to 0.0002 during the warmup period. AdamW was used as an optimizer. The CNN models were trained with the Adam optimizer, using learning rate of 0.001, no weight decay, and batch size of 32.

#### Code Repository

All code to derive results in this paper is available in a GitHub repository: https://github.com/ML-Bioinfo-CEITEC/genomic_benchmarks

## 3. Results

To evaluate the performance of the model on a testing set, we will use the F1 metric:

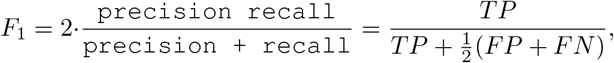

where TP is a number of true positives, FP a number of true positives and TN is a number of false negatives.

The running time has been 5-30 minutes for one run of CNN model and 2-6 hours for transformer models (Google Clound Platform, n1-highmem-8 virtual machine, NVIDIA Tesla T4 GPU).

The performance of the models summarized in F1 metric on a testing set is reported in the Table 2. As you can see DNABert is superior in all five our benchmark datasets and the fine-tuned DNABert outperformed the model with the randomized weights in four out of five cases.

**Table 2.** F1-score on testing sets, best model in bold font.

| EVALUATION: DATA SET | CNN | DNABERT (RANDOMIZED WEIGHTS) | DNABERT (PRETRAINED) |
| --- | --- | --- | --- |
| HUMAN_NONTATA_PROMOTERS | 83.9 ± 1.5 | 88.0 ± 0.7 | **91.9 ± 0.9** |
| HUMAN_OR_WORM | 83.3 ± 0.6 | 80.0 ± 0.6 | **96.8 ± 0.2** |
| HUMAN_ENHANCERS_ENSEMBL | 61.9 ± 1.4 | 83.8 ± 0.7 | **86.9 ± 0.3** |
| CODING_VS_INTERGENOMIC_SEQS | 74.8 ± 0.5 | 78.3 ± 0.6 | **92.8 ± 0.6** |
| HUMAN_ENHANCERS_COHN | 67.1 ± 0.6 | **76.5 ± 0.5** | 74.1 ± 0.4 |

## 4. Discussion

In this paper, we have experimented with transformers applied to classification of DNA sequences. We have shown that the model pre-trained on human genome achieves better accuracy than the same model with randomized weights and a convolutional neural network model. While ML researchers in the genomic field currently uses rather simple architectures like LSTMs and CNNs, the HuggingFace implementation of DNABert should encourage wider adoption of transformers.

DNA sequences present a unique challenge for machine learning because of their length and complexity. Transformers provide a more effective way to model these sequences than traditional neural networks. However, with only one transformer model trained over DNA available, many questions remain open for further investigation. Would the bigger models achieve better performance as for natural language and also for protein sequences (Rives et al., 2021), (Elnaggar et al., 2020), (Xiao et al., 2021)? If one universal DNA language model sufficient or would it be better to train a separate language model for each model organism (human, mouse, zebrafish, …).

And finally, taking into account the heterogeneous nature of human genome, would it be better to train on corpus that would not be the whole genome but rather a handcrafted specific subsample, e.g. for promoters taking only segments close to transcription splice site? This should be investigated in future work.

## Acknowledgement

P. Simecek’s work on this paper has been supported by H2020 MSCA IF grant LangOfDNA (Grant ID 896172).

